# T-cell cellular stress and reticulocyte signatures, but not loss of naïve T lymphocytes, characterize severe COVID-19 in older adults

**DOI:** 10.1101/2022.12.21.521463

**Authors:** Mladen Jergović, Makiko Watanabe, Ruchika Bhat, Christopher P. Coplen, Sandip A. Sonar, Rachel Wong, Yvonne Castaneda, Lisa Davidson, Mrinalini Kala, Rachel C. Wilson, Homer L. Twigg, Kenneth Knox, Heidi E. Erickson, Craig C. Weinkauf, Christian Bime, Billie A. Bixby, Sairam Parthasarathy, Jarrod M. Mosier, Bonnie J. LaFleur, Deepta Bhattacharya, Janko Ž. Nikolich

## Abstract

In children and younger adults up to 39 years of age, SARS-CoV-2 usually elicits mild symptoms that resemble the common cold. Disease severity increases with age starting at 30 and reaches astounding mortality rates that are ~330 fold higher in persons above 85 years of age compared to those 18-39 years old.

To understand age-specific immune pathobiology of COVID-19 we have analyzed soluble mediators, cellular phenotypes, and transcriptome from over 80 COVID-19 patients of varying ages and disease severity, carefully controlling for age as a variable. We found that reticulocyte numbers and peripheral blood transcriptional signatures robustly correlated with disease severity. By contrast, decreased numbers and proportion of naïve T-cells, reported previously as a COVID-19 severity risk factor, were found to be general features of aging and not of COVID-19 severity, as they readily occurred in older participants experiencing only mild or no disease at all. Single-cell transcriptional signatures across age and severity groups showed that severe but not moderate/mild COVID-19 causes cell stress response in different T-cell populations, and some of that stress was unique to old severe participants, suggesting that in severe disease of older adults, these defenders of the organism may be disabled from performing immune protection. These findings shed new light on interactions between age and disease severity in COVID-19.

## Introduction

Severe acute respiratory syndrome coronavirus-2 (SARS-CoV-2) infection is asymptomatic, mild or moderate in up to 95% infected humans [1, 2], and severe in 5% of patients. Outcomes of COVID-19 are remarkably age-specific, with most of the severe cases and deaths occurring in older adults [3, 4]. Infection fatality rate increases in a log-linear fashion with age in individuals older than 30 years [4, 5], and culminates to be ~270-fold higher in those over 80 relative to the population 18-39 years old [6]. In addition to age, risk factors for severe disease include hypertension, obesity, type 2 diabetes and male sex [7]. Importantly, geriatric frailty assessed by a clinical scale is associated with COVID-19 severity even more closely than chronological age [8].

Correlates of disease severity were thoroughly studied. Severe cases display increased viral load and delayed viral clearance [9, 10] implying inadequate virus control by the immune system. In addition, multiple perturbations of both innate and adaptive immune parameters have been reported to predict disease severity [11–13]. Severe COVID-19 was reproducibly accompanied by elevated levels of inflammatory cytokines, namely interleukin-6 (IL-6), C-reactive protein (CRP), and Tumor Necrosis Factor-α (TNF-α) [14, 15]. Furthermore, multiple cellular immunity defects were associated with COVID-19 severity, including decreased lymphocyte and increased neutrophil blood counts with an inverted neutrophil to lymphocyte ratio as a strong predictor of mortality [16]. Decreased naïve CD4 T-cell counts were also linked to severe COVID-19 [17]. However, the major caveat affecting early cohort studies on severe COVID-19 was the lack of subject groups of comparable age experiencing moderate or mild disease [11, 12, 18, 19] as well as uninfected age-matched controls. Multiple studies have compared older severe subjects to moderate subjects of younger age, potentially confounding interpretation. Even with advanced age, most patients will not present with severe disease. Therefore, to fully understand immune pathology associated with severe disease, proper age matched controls are essential.

Here we analyzed blood samples from patients with confirmed COVID-19 across a spectrum of age and severity, including older subjects with moderate/mild disease. We measured levels of soluble factors (IL-6, CRP, TNF-α. IL-1β); immune cell phenotypes; levels of apoptosis; single cell RNA-Seq profiles and antigen specific T-cell responses. We discovered that increased reticulocytes, neutrophils, CRP and IL-6 strongly correlate with disease severity irrespective of age. In our study, however, decreased naïve T-cell numbers were measured in older patients experiencing moderate, severe or no disease alike, confirming our prior results [20] and suggesting that this is an age-associated, and not severity-associated phenotype. Single-cell RNA sequencing (scRNA-Seq) revealed a strong reticulocyte transcriptional signature selectively in severe COVID-19, and confirmed by increased circulating reticulocyte numbers, implying a critical role of oxygen deficit in severe disease. Other data from the scRNA-Seq revealed pathways consistent with general cellular stress and respiratory/metabolic crisis in immune cell subsets in severe COVID-19 of older participants.

## Methods

### Study participants

This study was approved by the University of Arizona IRB (protocol #1510182734 and #1410545697) and the Oregon Health and Science University IRB (protocol #00003007) and was conducted in accordance with all federal state and local regulations and guidelines. The cohort included 86 SARS-CoV-2 positive individuals recruited in Tucson and Phoenix (AZ) and Indianapolis (IN); confirmed positive by PCR test and 64 age matched healthy community-dwelling individuals recruited in Tucson (AZ) and Portland (OR) prior to September 2019. Serum was collected from additional 17 patients hospitalized with non-COVID-19 related ARDS in Tucson (AZ). Healthy participants received nominal participation incentive. All SARS-CoV-2 positive participants underwent venipuncture within 3 weeks from symptom onset (mean=11.57 ± 5.04 days). COVID-19 participants were scored for disease severity according to WHO clinical progression scale [21]. Study participants were divided into the following groups; moderate adult (MOD-A), moderate old (MOD-O), severe adult (SEV-A), severe old (SEV-O), adult healthy controls (CTRL-A) and old healthy controls (CTRL-O). 47% of SARS-CoV-2 positive participants were female, and healthy controls were age- and sex-matched. Blood was drawn into BD Vacutainer Blood Collection Tubes with Sodium Heparin (BD Bioscience, Franklin Lakes, NJ) and in BD Vacutainer tubes with K2EDTA (for complete blood counts). Peripheral blood mononuclear cells (PBMC) were separated by Ficoll gradient separation from heparinized blood and cryopreserved in fetal calf serum (FCS) + 10% DMSO. Cryopreserved PBMC (5 x 10^6^/sample) were thawed in prewarmed RPMI-1640 media supplemented with L-glutamine (Lonza, Basel, Switzerland) + 10% FCS. Thawed PBMCS were rested overnight at 37 °C in X-VIVO^™^-15 Medium (Lonza) supplemented with 5% human-AB serum.

### Cytokine measurements

Plasma concentrations of cytokines IL-6, CRP, TNF-α and IL-1β were measured by Milliplex Map Magnetic Bead Assays (MilliporeSigma, Burlington, MA; IL6, IL-1β, TNF-α multiplex assay; CRP assay) and acquired on a Magpix instrument (Luminex, Austin, TX), per manufacturer instructions.

### Flow cytometry

Cells were stained with surface antibodies in PBS (Lonza) + 2% FCS, stained with live/dead fixable blue dye (Thermofisher, Waltham, MA) and then fixed and permeabilized using FoxP3 Fix/Perm kit (eBioscience, San Diego, CA). Samples were acquired using a Cytek Aurora cytometer (Cytek, Fremont, CA) and analyzed by FlowJo software (Tree Star, Ashland, OR). Flow cytometry files were uploaded to Cytobank, a cloud-based computational platform where Citrus clustering was performed on live singleT-cells in the lymphocyte gate. We ran Nearest Shrunken Centroid (R implementation of Prediction Analysis for Microarrays (PAMR)) – a predictive association model within Citrus, with minimal cluster size of 2% of all cells and a false discovery rate of 1% for highest stringency. A list of antibodies is available in the Supplemental Table 1.

### Single cell RNA sequencing

PBMC samples from 15 COVID-19 patients (n=3-4/age and severity group) were used to generate single cell RNA seq libraries using the 10x genomics Chromium Single Cell 3’ Library & Gel Bead Kit v3 and Chromium Single Cell V(D)J Reagents kit as per manufacturer’s protocols. The overall sequence depth was ~ 50000 reads/cell using the NovaSeq 6000 platform (Illumina, San Diego, CA). The demultiplexed reads obtained from NovaSeq were first aligned to a reference human genome assembly (GRCh38-3.0.0) using the Cell Ranger pipeline (10X Genomics, Pleasanton, CA). The feature barcode matrices, overall sample summaries and cloupe & vloupe files were generated using the Cell Ranger count. Further downstream analysis was done using Seurat R package v4.0 [22–25]. QC analysis was performed on each individual feature barcode matrix. All the cells expressing less than 200 genes were discarded, we filtered out cells expressing >10% of mitochondrial genes and performed doublet scoring and removal using the ‘scrublet’ package [25]. Normalization, scaling, dimensionality reduction, clustering, visualization, reference-based cell type labelling was performed using Seurat and ggplot2 [26]. Cell type-wise differentially expressed genes for different categories were obtained using the ‘Seurat::FindMarkers’ function with test method as MAST [27]. Pathway and process enrichment analysis were done via Gene Set Enrichment Analysis (GSEA) using GSEA v4.2.3 [28] software with the gene set “c5.go.v7.4.symbols.gmt [Gene Ontology]” of MSigDB [29] and keeping the permutations random sampling as 1000 (default). The false discovery rate (FDR) <0.1 and the normalized enrichment score (NES) were utilized to identify the significant enrichment pathways.

### ELISpot assays

T-cells specific immunity to peptide mix corresponding to Spike, Nucleocapsid and Matrix proteins were measured by ELISpot detection of IFN-γ. PBMCs were stimulated in X-VIVO 15 media with 5% male human AB serum containing either 0.6 nmol PepTivator SARS-CoV-2 Prot S, N and M peptide pool (Miltenyi Biotec, Bergisch Gladbach, Germany) for antigen specific T-cell stimulation, or positive control (anti-CD3 mAb) or blank media as negative control. Cell suspensions were transferred to pre-coated Human IFN-γ ELISpot plus kit plates (Mabtech, Nacka Strand, Sweden) and developed after 20h per manufacturer’s instructions. Spots were quantifiedn using an ImmunoSpot Versa (Cellular Technology Limited, Cleveland, OH).

### Statistical analysis

SPSS, Graph Pad Prism, and the R statistical packages were used for analysis. Group differences were calculated either by Mann Whitney U-test, one-way ANOVA or Kruskal Wallis test with post hoc correction based on model assumptions (normality of residuals, heterogeneity of variances, etc). We used multivariable linear regression to predict both severity and days from onset of symptoms. Missing data were imputed using multiple imputation and predictive mean matching through the Hmisc package in R [30]. Graphical representation of the contribution of each predictor variable to the fitted model after adjusting for all other predictors is shown by graphical representation of an ordered dot plot for each predictor (y-axis) by the partial χ2 test statistic scaled by the degrees of freedom for each variable (x-axis); the partial χ2 test statistic and p-values are shown on the left side of the graphs. In addition to showing relative contribution of each of the predictors, we use R^2^ as a metric of predictive capacity of the model. With a sample size of 99 and 27 potential predictor variables there was a high degree of potential overfitting (e.g., overoptimization of model performance). We calculated overfitting by comparing R^2^ across 200 bootstrap resamples and reported a corrected R^2^ as the difference between the original R^2^ and the average R^2^ across the 200 bootstrap resamples. Additionally, to further reduce model overfitting, variable reduction was performed by penalized maximum likelihood described in Friedman et al [31]. For all statistical differences *p<0.05, **p<0.01, ***p<0.001. ****p<0.0001.

## Results

### Study participant demographics and disease severity

We enrolled 86 subjects with SARS-CoV-2 infection confirmed by PCR and compared to controls sampled prior to September 2019. We have stratified participants into groups matched by disease severity and age (**Supplemental Figure 1A**). We scored disease severity across a range from 0 (not infected) to 10 (deceased) according to the WHO Clinical Progression Scale[21] with scores ≥6 (receiving oxygen by non-invasive ventilation) considered severe (**Supplemental Figure 1B**). If the subject progressed to a worse clinical outcome the highest recorded score was used. COVID-19 disease severity and infection fatality risk increases sharply after age 50 [32, 33]. Therefore, we classified all the participants less than 50 years old (yo) as adults, and those over 55 as older (subjects 50-55 yo were excluded from the study). By these standards our total group size was N=18 adults with moderate disease (MOD-A, average age 34.17 ± 7.64), N=22 older adults with moderate disease (MOD-O, age 74.23 ± 9.42), N=24 adults with severe disease (SEV-A, age 39.67 ± 9.46) and N=22 (SEV-O, age 74.50 ± 6.91) older adults with severe disease. All subjects were sampled within 3 weeks of symptom onset and groups did not differ in time from onset (**Supplemental Figure 1C**). In addition, we obtained serum samples from 17 patients hospitalized with non-COVID-19 related ARDS who were 38-82 years old (average 56.11 ± 11.85 years).

### Neutrophil: lymphocyte ratios, IL-6 and CRP, but not TNFα, predict severe COVID-19 regardless of age

We analyzed blood leukocyte counts in severe and moderate subjects stratified by age. We found that neutrophils were increased in both COVID-19 SEV-A and SEV-O, as well as non-COVID ARDS (**Figure 1A**). On the other hand, lymphopenia was pronounced only in older adults with severe disease (**Figure 1B**). Median lymphocyte count in SEV-A was reduced compared to CTRL-A (0.99 k/μl vs. 1.75 k/μl) but the difference was not statistically significant (**Figure 1B**). For neutrophil to lymphocytes ratios, both COVID-19 severe groups and non-COVID ARDS exhibited higher values compared to age-matched controls (**Figure 1C**). We found no difference between the groups in monocyte (**Figure 1D**) or eosinophil counts (**Figure 1E**). Basophils were decreased in both severe COVID-19 groups and non-COVID ARDS (**Figure 1F**). Some of the cytokines previously reported to be increased in severe COVID-19 like IL-6[34], CRP and TNF-α [35] are known to be increased with age even in the absence of infection. We show that increased levels of IL-6 (**Figure 1G**) and CRP (**Figure 1H**) are associated with severe COVID-19 regardless of age as well as of non-COVID ARDS. On the other hand, we found TNF-α to be increased only in SEV-A but not SEV-O (**Figure 2I**), with no difference between the groups in levels of IL-1β (**Figure 2J**). Therefore, irrespective of age, severe COVID-19 remains characterized by increased neutrophil to lymphocyte ratio, elevated CRP and IL-6, the features shared by non-COVID ARDS, whereas TNFα elevation is found in SEV-A and non-COVID ARDS, but not SEV-O COVID-19.

**Figure 1.**
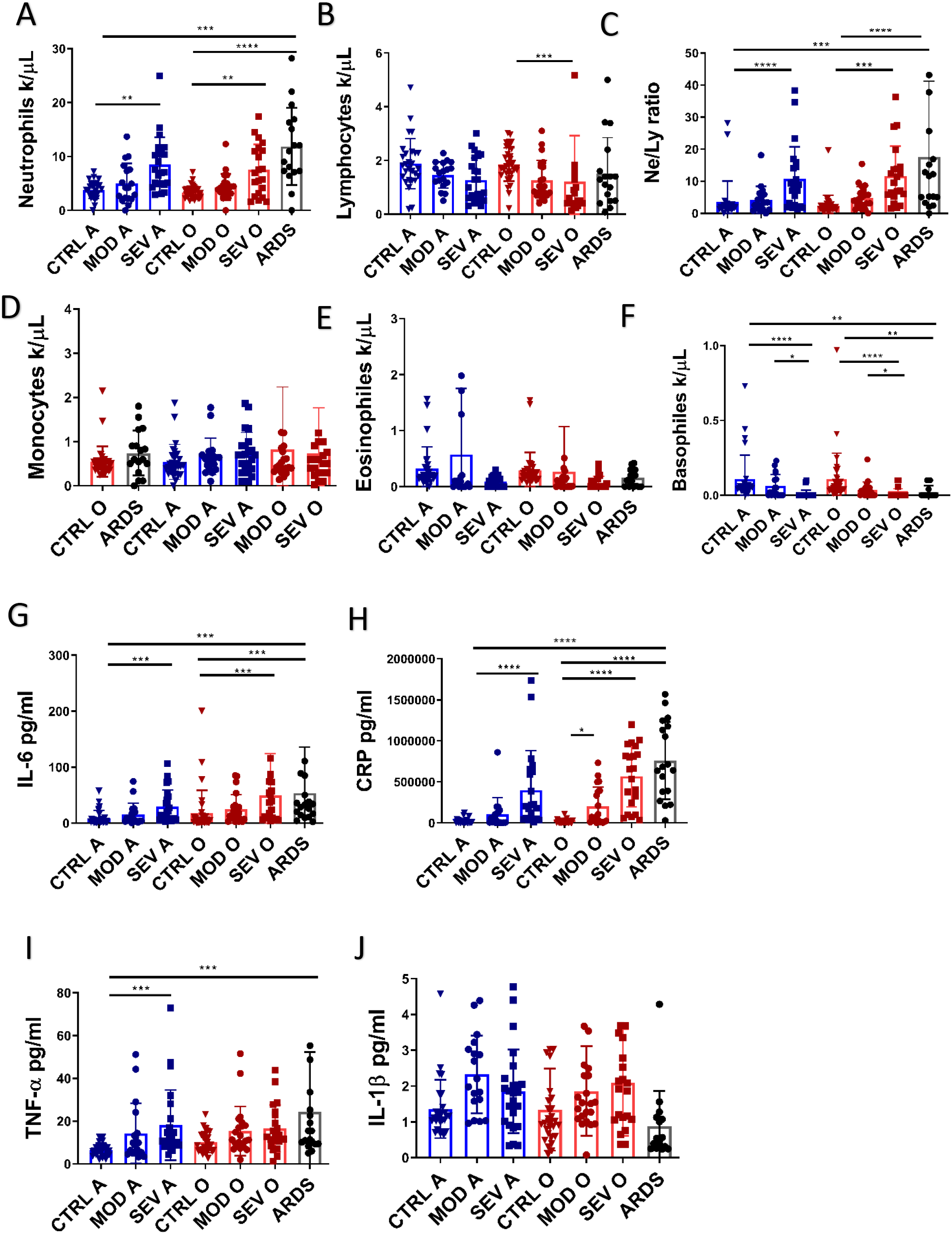
Complete blood counts (CBC) were obtained from fresh blood of all participants **A)** neutrophil counts were severe COVID-19 as well as non-COVID ARDS, **B)** Decreased lymphocyte counts were statistically significant only in older adults with severe COVID-19, **C)** both COVID-19 severe groups and non-COVID ARDS had increased neutrophil to lymphocytes ratios. Monocyte **D)** and Eosinophil **E)** counts did not differ among the groups. **F)** Basophils were decreased in both severe COVID-19 groups and non-COVID ARDS. **G)** IL-6 serum levels were elevated both severe COVID-19 groups and non-COVID ARDS as well as **H)** C-reactive protein (CRP). **I)** TNF-α was increased only in adults with severe COVID-19. **J)** No difference between the groups in levels of IL-1β. Data presented as mean ± standard deviation. Kruskal-Wallis test with Dunn’s post hoc correction. For all statistical differences *p<0.05, **p<0.01, ***p<0.001. ****p<0.0001.

**Figure 2.**
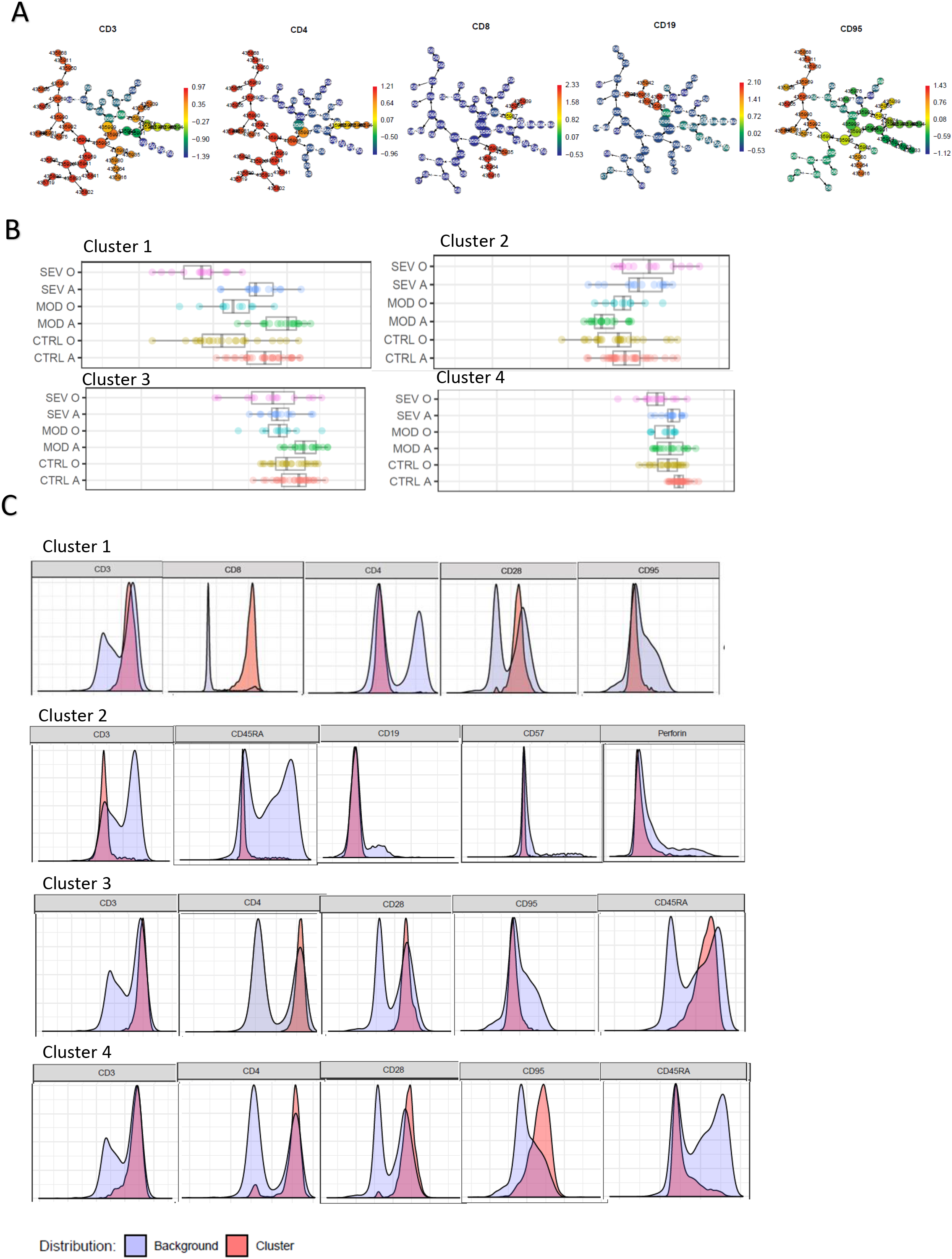
Nearest Shrunken Centroid (PAMR) – a predictive association model within Citrus, with minimal cluster size of 2% was applied to 32 parameter flow cytometric panel. **A)** Expression of common phenotyping markers on identified cell clusters, **B)** four cell clusters were differentially expressed among the groups. **C)** phenotypic markers identify lineage of cells in clusters 1-4

### COVID-19 severity analysis with age by unbiased clustering of flow cytometry data

We designed a 32-marker flow cytometric panel to analyze PBMCs from COVID-19 subjects and healthy controls (antibody list in **Supplemental Table 1**. Given the high dimensionality of our flow cytometric panel, we first applied an unbiased computational approach for subpopulation clustering by using Cytobank Citrus to identify cell subsets differentially represented between the groups (**Figure 2A**). The pamr package in R was used to classify abundance clusters necessary to differentiate the groups, resulting in four clusters (**Figure 2B**). Cluster 1 exhibited a CD3+CD8+CD28+CD95-phenotype, corresponding to naïve CD8 T-cells, and this cluster was less abundant in all three older groups (uninfected control, moderate and severe) compared to their adult counterparts **(Figure 2C)**. Cluster 2 did not express any of the markers in the staining panel (**Figure 2C**) and was elevated in both SEV-A and SEV-O compared to all other groups. Cluster 3 exhibited a CD3+CD4+CD28+CD95-CD45RA+ phenotype, representing naïve CD4 T-cells and was decreased in SEV-A, SEV-O and MOD-O. Cluster 4 was CD3+CD4+CD28+CD95-CD45RA-corresponding to central memory CD4 T-cells and was decreased in SEV-O compared to all the other groups. Thus, overall unbiased clustering identified several T-cell subpopulations which differed in abundance between the groups and a cluster of unknown origin which was elevated in both severe groups irrespective of age.

### Reticulocyte accumulation is a robust and age-independent predictor of COVID-19 severity

To identify the nature of the cluster 2 cells, which was elevated in severe subjects irrespective of age, we performed single-cell RNA sequencing (scRNA-seq) on 15 COVID patients from each group to gain insight into transcriptional signatures and pathways associated with severity. **Table 1** shows the demographics and the number of cells analyzed individually for each patient. By mapping our samples to the Seurat package reference dataset and using unsupervised clustering (see Methods), we identified 30 cell types across all samples (**Figure 3A, Supplemental Figure 2A**). The scRNA data was classified by severity and age groups (**Table 1**)’ with the composite UMAP color-coded by classification shown in **Supplemental Figure 2B**. The expression of “HBB”, a component of hemoglobin, was observed to be higher in severe patients as compared to the mild ones (**Figure 3B**). Red blood cells are separated from the PBMCs by ficoll separation. We thus hypothesized that nucleated cells expressing hemoglobin genes might be reticulocytes, potentially generated in response to hypoxia and anemia in severe COVID-19 cases.

**Figure 3.**
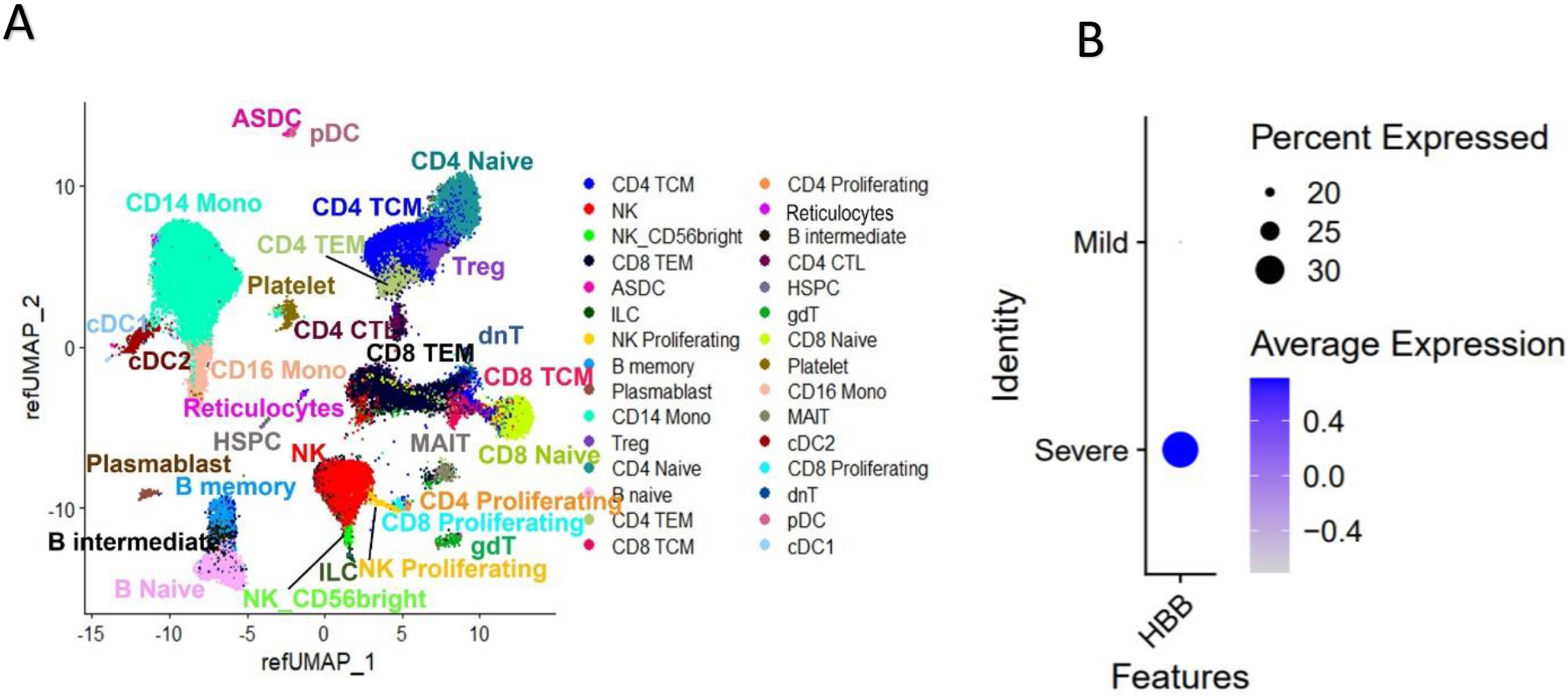
**A)** UMAP projections of 92715 cells captured from 15 COVID patients annotated by cell types. **B)** Dot plot of HBB expression levels and percentage of in severe and mild patients (6 severe and 9 mild).

**Table 1.**
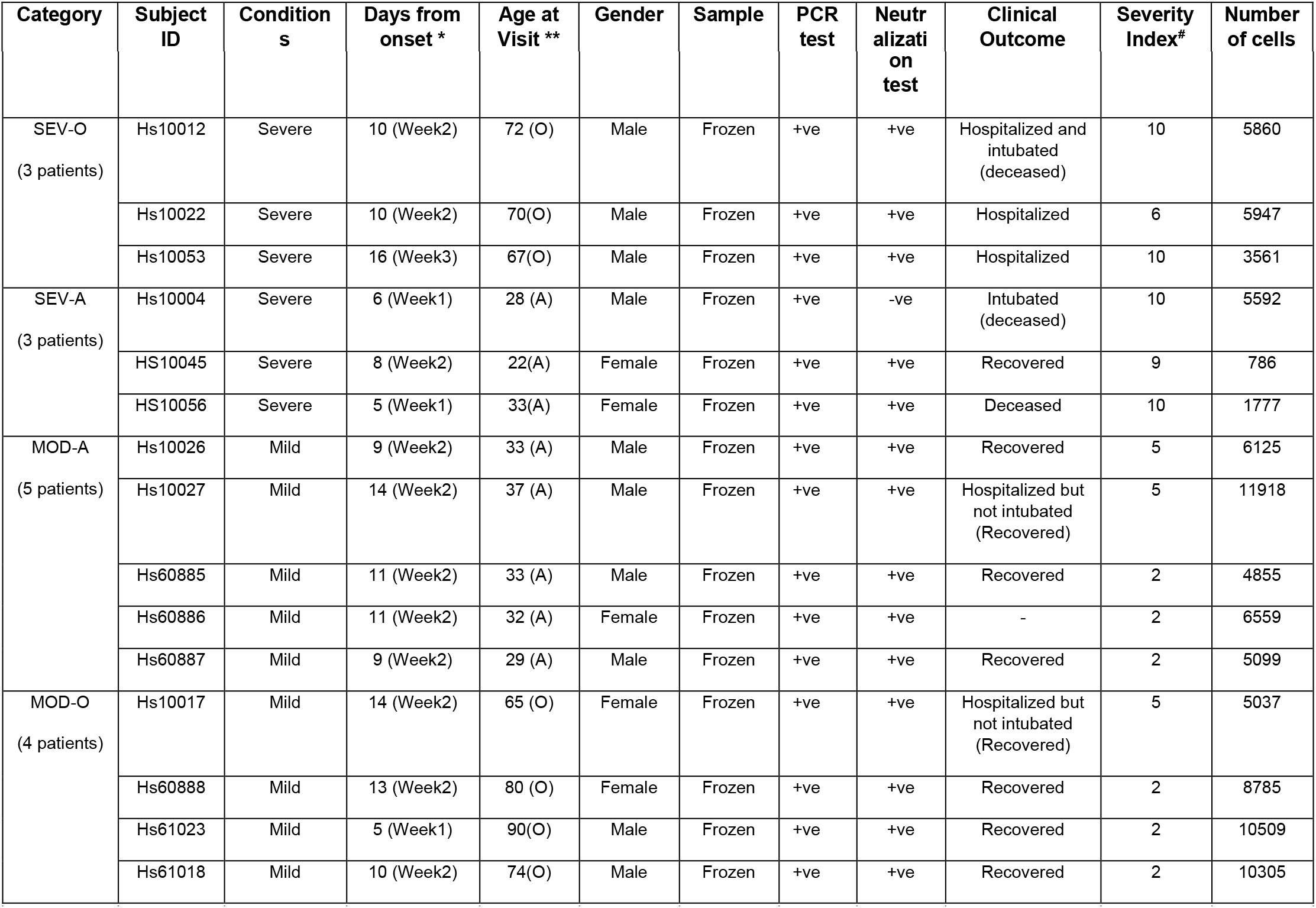
Demographics, sample characteristics and disease course data for the 15 COVID-19 patients. *PBMCs were collected between 5 to 16 days from the onset of symptoms as shown in column ‘Days from onset’. **Patients with age < 65 years were considered Adult (A) and age ≥ 65 years were considered as Old (O); out of 15 patients 7 were old and 8 were adult. #The severity Index is calculated based on clinical outcome using WHO defined severity score (doi:10.1016/ S1473-3099(20)30483-7).

Reticulocytes express the transferrin receptor (CD71) and do not express leukocyte common antigen CD45 [36]. We therefore performed additional flow cytometric staining for these markers in all participants. We confirmed the existence of this CD45-CD71+ population within the lymphocyte gate selectively in severe cases (**Figure 4A**). Given that this population was contained within the lymphocyte gate by forward and side scatter, we excluded the possibility that these are mature erythrocytes. Furthermore, this population did not express any of the classical lineage markers of immune cells such as CD3, CD19, CD16, CD11b and CD15 but did express higher levels of CD147 (Basigin) **(Figure 4B)**, known to be highly expressed on reticulocytes [36]. This cell population was elevated in both severe groups irrespective of age compared to all the other groups (**Figure 4C**).

**Figure 4.**
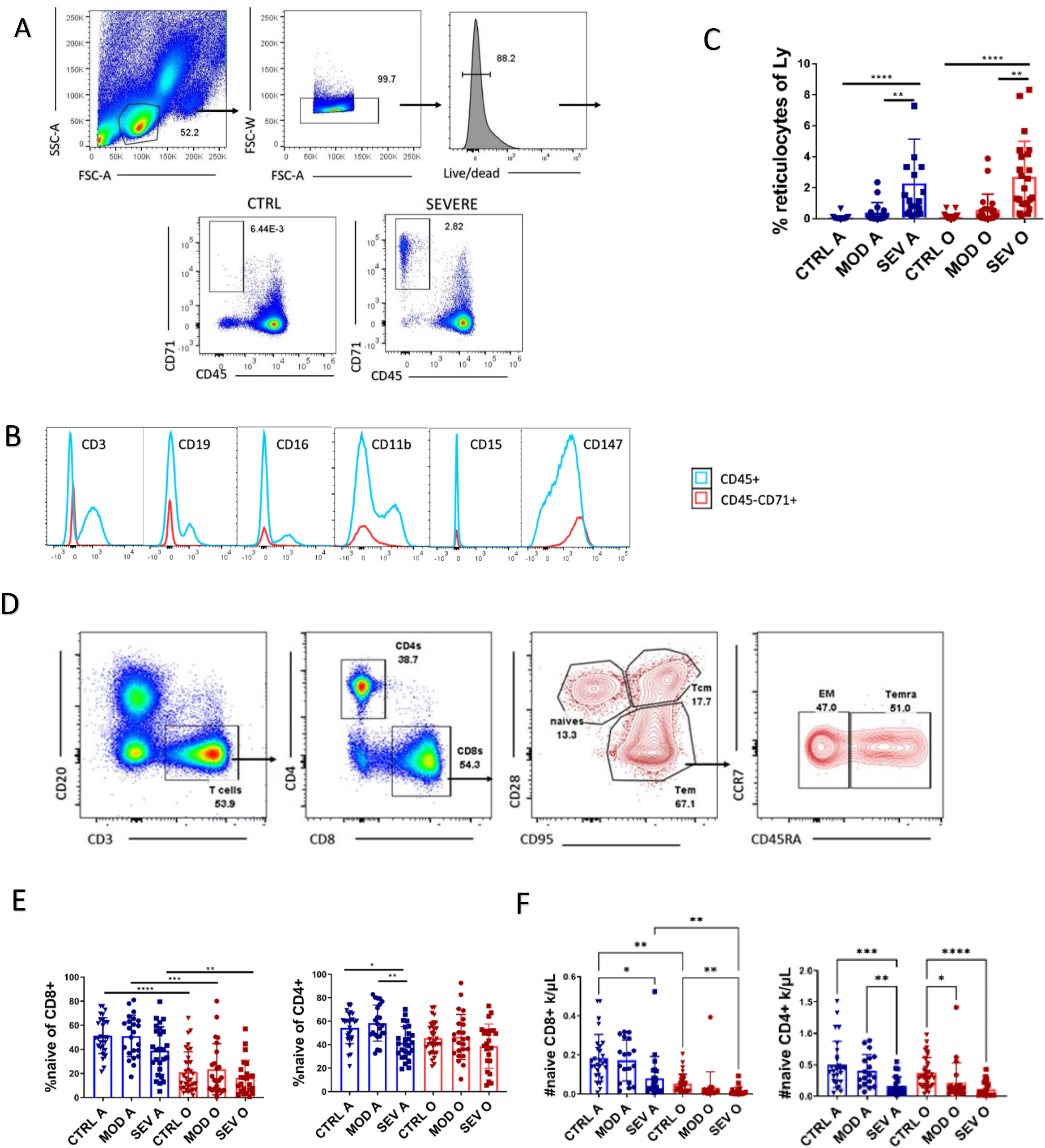
**A)** Flow cytometric gating strategy used to identify CD45-CD71+ cells. **B)** No expression of CD3, CD19, CD16, CD11b and CD15 but high expression of CD147 on CD45-CD71+ cells **C)** Percentage of CD45-CD71+ cells within the lymphocyte gate was increased in subjects with severe COVID-19. **D)** Flow cytometric gating strategy used to identify T cell subsets. **E)** Decreased proportion of naïve T-cells was a characteristic of all three advanced age groups irrespective of SARS-CoV-2 status **F)** Absolute counts of naïve T-cells were lower in severe COVID-19 compared to age matched controls. Data presented as mean ± standard deviation. Kruskal-Wallis test with Dunn’s post hoc correction. For all statistical differences *p<0.05, **p<0.01, ***p<0.001. ****p<0.0001.

### Naïve CD8 T-cell loss is an age-sensitive trait not linked to COVID-19 severity

To further confirm Citrus clustering, we performed traditional manual gating of T-cell phenotypes **(Figure 4D).** Similar to Citrus clustering, flow cytometric gating showed that decreased proportion of naïve CD8 T-cells was a characteristic of all three advanced age groups (CTRL-O, MOD-O and SEV-O) and not associated with COVID-19 severity (**Figure 4E**). When translated into absolute counts per μL of blood, due to overall lower lymphocyte counts, the severe cases did have lower naïve T-cell counts than their age-matched controls (**Figure 4F**). However, older adult controls also had lower naïve CD8 counts compared to their adult counterparts. Moreover, naïve CD4s were also decreased in MOD-O, thus we cannot conclude that a decrease in naïve T-cells is a general feature of severity. Other T-cell subsets, such as central memory T-cells, were decreased in absolute count in severe cases **(Supplemental Figures 3A and 3B**) although not decreased as a proportion of total T-cells. The proportion of CD8+ T-cells with effector/memory (Tem) phenotype was increased in severe adults (**Supplemental Figure 3C**), but there was no difference in proportion of terminally differentiated Temra cells. In both cases this did not translate into any difference in absolute numbers between the groups (**Supplemental Figure 3D**). Additionally, we measured antigen-specific T-cells reactive to overlapping peptide pools spanning the entire Spike (S), nucleocapsid (N) and membrane (M) by ELISPOT. There was no difference in number of N, M, or S antigen-specific T-cells between SARS-CoV-2 positive groups by age or severity (**Supplemental Figure 3E**). Moreover, we observed very few cross-reactive T-cells in samples from control participants recruited prior to the pandemic (**Supplemental Figure 3E**).

### Reticulocytes, CRP levels and neutrophil count are the variables most predictive of COVID-19 severity

To integrate and directly compare severity-related parameters for their predictive strength, we analyzed how all cytokine and cell flow cytometry variables correlated to patient severity scores and time from disease onset as a potential confounder. Variable selection was performed using regularized regression and selected variables were visualized by ranking the “importance” based on Wald χ^2^ test statistics. We calculate overfitting by comparing R^2^ across 200 bootstrap resamples and report a corrected R^2^ as the difference between the original R^2^ and the average R^2^, this measure can be used as an internal validation. Bootstrap resampling on all 27 predictor variables resulted in poor model performance for both outcome variables. RBC and CRP displayed the highest adjusted Wald test statistics for severity (**Supplemental Figure 4A**) and percentage of CD8+ and CD4+ T-cells for days since onset of COVID-10 symptoms (**Supplemental Figure 4B**); For both severity and time from onset there were only two variables that were statistically significant when all measured variables were included. Therefore, we used regularized regression to reduce the number of variables and comparisons; selected variables included percent reticulocytes; Ba; RBC; % TEM of CD8; % TEMRA; TNFα; IL6; and CRP for the severity outcome. For the days from COVID-19 symptom onset Mo; Eo, Ba; and IL6 were selected. The severity outcome resulted in 6 statistically significant variables with p-values < 0.05 and a corrected R^2^ of 0.575, with reticulocytes, CRP and neutrophil counts were most predictive of disease severity (**Figure 5A**). Days from symptom onset was not associated with any potential predictors, and also demonstrated by a poor predictive fit (R^2^ = 0.20) (**Figure 5B**). We assume that observed differences were not influenced by individual differences in time from disease onset between COVID19 groups, with means (SD) in the MOD-A, MOD-O, SEV-A and SEV-O of 13.5 (5.9); 13.11 (7.1); 14.2 (6.9); and 11.9 (6.5), respectfully. Thus, we conclude that in all severe subjects, irrespective of age, reticulocytes, CRP levels and neutrophil counts were the variables most robustly associated with COVID-19 severity scores.

**Figure 5.**
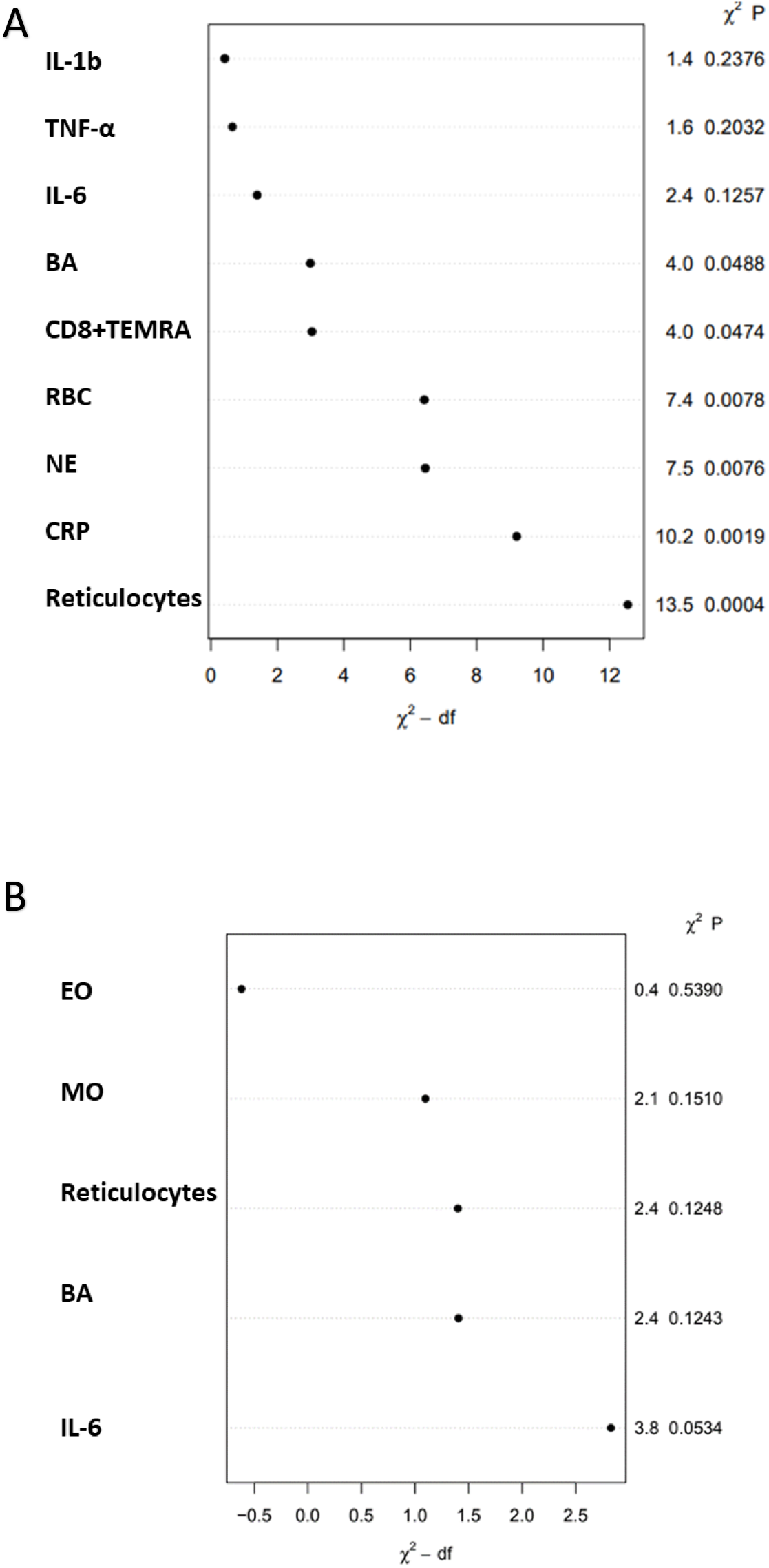
Multivariable linear regression to predict dependence of **A)** severity and **B)** days from onset of symptoms on measured cell and serum variables. Predictor variables were selected using regularized regression and the elastic-net algorithm.

### Cellular stress and crisis signatures characterize transcriptomes of multiple T-cell subsets in older participants with severe COVID-19 (SEV-O)

To begin to address the condition- and age-wise transcriptional differences across the samples, we performed additional scRNA-seq analysis on a subset of patients (described in Table 1) and applied differential expression analysis genes between the pairwise categories such as SEV-O and MOD-O (Methods). Differentially expressed genes with an adjusted p-value (adj. p-val) □ 00001 obtained from such comparisons, along with their log2 fold change (logFC) values were subjected to pathway analysis. GSEA was used to identify the pathways that were enriched for either severity or for age. Individual pathway enrichment comparisons are shown in Figure 8 for selected CD4 and CD8 T-cell subsets comparing SEV-O to MOD-O and SEV-O to SEV-A groups. We found consistent enrichment of cellular and oxidative stress, catabolism, starvation and negative regulation of biosynthetic processes (including RNA metabolism) pathways in five T-cell subsets of SEV-O patients: CD8 naïve (Supplemental **Figure 5A**) and effector memory (**Supplemental Figure 5B and 5C**), and CD4 central memory **(Supplemental Figure 5D)**, effector memory **(Supplemental Figure 5E and 5F**) and cytotoxic (CTL) clusters **(Supplemental Figure 5G and 5H)**. By contrast, in these same cell subsets MOD-O and SEV-A participants exhibited enrichment for antiviral immune pathways, including negative regulation of viral processes, upregulation of interferon response and cytokine responses, and defense responses to microorganisms. While these results require detailed experimental substantiation, overall pathway enrichment suggests that while COVID-19 transcriptional profiles in T-cells of MOD-O and even SEV-A patients involved immune defense pathway activation across T-cell subsets, their SEV-O counterparts were unable to mount such defense and were instead engaged in cell survival in the face of severe cellular stress.

### CD8 but not CD4 T-cell apoptosis characterizes COVID-19, and naïve CD8 T-cell apoptosis selectively predicts disease severity

Significantly enriched pathways related to cellular stress, catabolism and death prompted us to examine whether COVID-19 severity also mechanistically correlated to lymphocyte death. We plotted the expression of caspase 3 (CASP3), a member of the final effector enzyme complex that mediates apoptotic cell death [37, 38] within the scRNA-seq data (**Figure 6A)** and found its expression to be even across multiple cell types. We assessed functional importance of these observations by measuring Annexin V and CASP-3 staining by flow cytometry. Annexin V is commonly used for staining of apoptotic cells due to its ability to bind phosphatidylserine which is translocated on the outer membrane in apoptotic cells while caspase-3 is a part of apoptosis signaling in T-cells [37, 38] (see gating strategy for flow cytometric analysis of T-cell apoptosis in **Figure 6B**. We found that caspase-3 was significantly elevated in CD8+ T-cells but not CD4+ T-cells selectively in SEV-O T-cells **(Figure 6C)**, whereas Annexin-V staining showed a similar trend but was not statistically significant (**Figure 6D**). Upon closer examination of CD8+ T-cell subsets it was revealed that caspase-3 was elevated only in naïve and Tcm CD8+ T-cells but not Tem **(Figure 6E)**, suggesting that severe COVID-19 may result in bystander T-cell apoptosis. Pathway enrichment analysis showed a positive correlation (FDR q-value <0.1) for intrinsic apoptotic signaling pathway supporting the above observation (**Supplemental Figure 6A**). Top genes from the leading edge for this pathway clearly show higher expression in SEV-O participants as compared to other participants (**Supplemental Figure 6B**).

**Figure 6.**
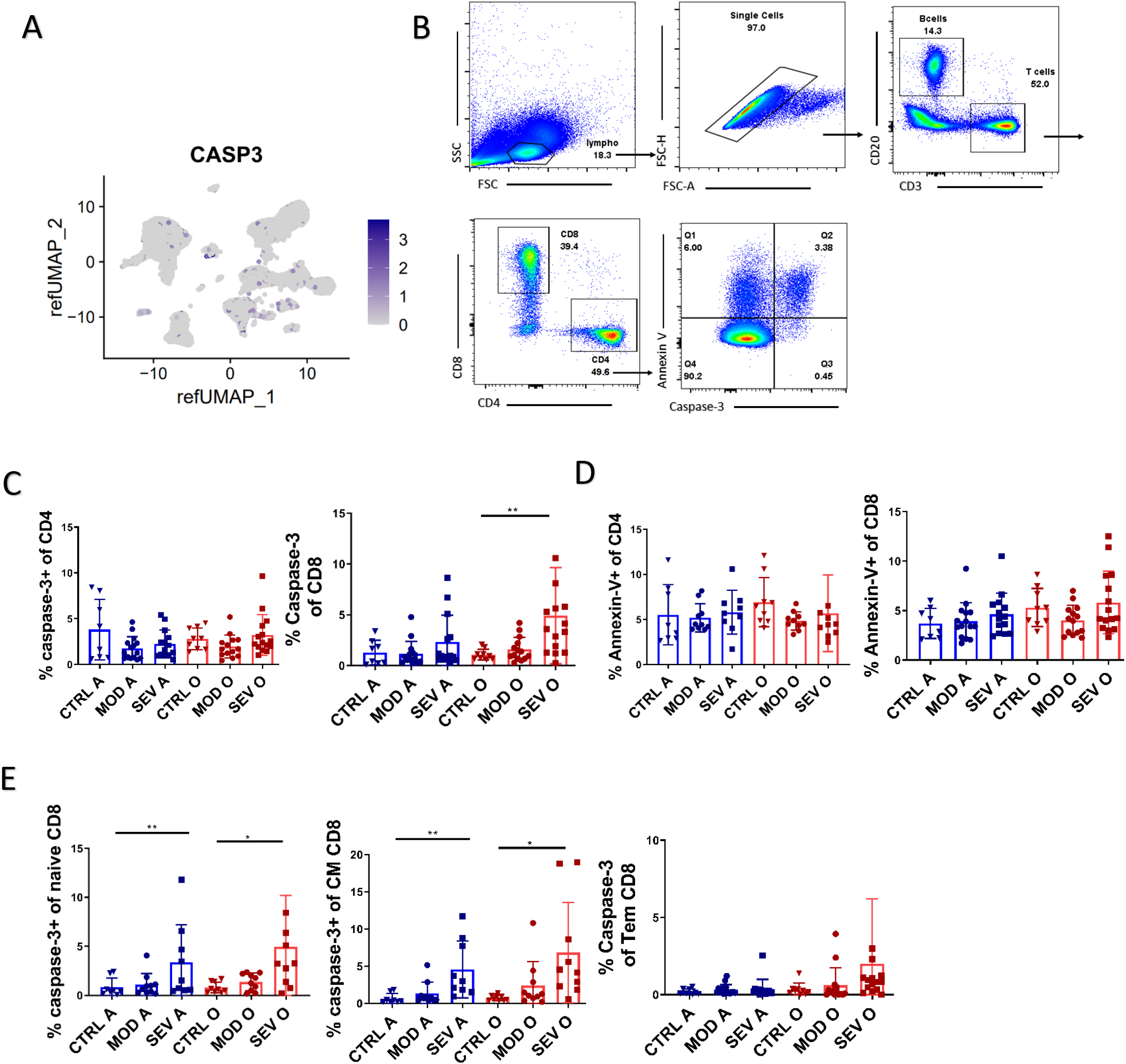
**A)** UMAP plot showing expression of CASP3 gene to be distributed evenly across all cells (cell type localization on the same UMAP is provided in Figure 4A). **B)** Representative flow cytometric gating to identify Annexin-V and Caspase-3 positive T-cells. **C)** Increased percentage of caspase-3 positive CD8+ T-cells in older adult severe group **D)** No difference in Annexin-V expression between the groups. **E)** Caspase-3 was increase on naïve and central memory CD8+ T-cells in both severe groups. Kruskal-Wallis test with Dunn’s post hoc correction. Data presented as mean ± standard deviation. For all statistical differences*p<0.05, **p<0.01, ***p<0.001. ****p<0.0001.

## Discussion

COVID-19 severity is associated with impaired viral control, as severe cases have been found to have higher viral titers and delayed viral clearance[9], uncoordinated adaptive immune responses [17] and hyperinflammation [39]. To gain further insight into immune phenotypes associated with COVID-19 severity we recruited a cohort which included adult and older demographic groups with severe COVID-19 and age matched controls with moderate disease. In many previous studies participants with severe COVID-19 were on average much older than participants with moderate disease, potentially confounding interpretation of results. We show that decreased proportion and absolute number of naïve T-cells are manifested by participants with severe COVID-19 but also by older adults with moderate disease and even older adult healthy controls. On the other hand, we show that neutrophilia and increased neutrophil/lymphocyte ratio is characteristic for all severe subjects irrespective age. This was not surprising, as abundant evidence implicated neutrophils and increased neutrophil extracellular trap formation as playing a central role in coagulopathy, organ damage, and immunothrombosis in severe COVID-19 [40–42]. Elevated IL-6 and CRP were characteristic of severe COVID-19 irrespective of age and were elevated to similar levels in non-COVID-19 ARDS.

Multivariable regression showed that increased reticulocytes, CRP levels and IL-6 were most predictive of COVID severity score. To our knowledge this is the first report documenting association of increased reticulocytes with COVID-19 severity score. One previous study showed increased reticulocyte counts in critically ill hospitalized patients compared to non-critical [43]. However, multiple studies have shown that anemia is associated with severe illness in COVID-19 [44, 45]. Thus, an increase in erythrocyte precursors can be viewed as an adaptation to anemia. scRNA-Seq confirmed the reticulocyte signature and revealed signaling pathways associated with cell stress and apoptosis were elevated in severe COVID-19, which was then confirmed by apoptosis assays, documenting selective upregulation particularly in old participants.

Overall, our results document new biomarkers of COVID-19 severity (reticulocytes) and further confirm the value of some (CRP, IL-6, neutrophil:lymphocyte ratios) biomarkers of COVID-19 severity as being universally applicable regardless of age, while refuting predictive values of others (numbers of Tn CD8 cells, which are solely sensitive to age). Finally, our results also contribute to mechanistic understanding of severe COVID-19 by identification of cellular stress and apoptotic responses in the very cells that are supposed to be defending the organism against the virus.

## Acknowledgments

Supported in part by the USPHS awards from the National Institutes of Health, National Institute on Aging, R37AG020719 and the Bowman Professorship in Medical Sciences to J.N-Ž. This project was supported, in part, with support from the Indiana Biobank and the Indiana Clinical and Translational Sciences Institute funded, in part by Grant Number UL1 TR002529 from the National Institutes of Health, National Center for Advancing Translational Sciences, Clinical and Translational Sciences Award and the National Center for Research Resources, Construction grant number RR020128, the Lilly Endowment and U01 – AG060900 from NIA. We thank contributors who collected samples and/or data used in this study, as well as subjects whose help and participation made this work possible. We would also like to thank Anastasia Kousa from Marcel van den Brink Lab for her valuable suggestions during preparation of the manuscript

## Author Contributions

M.J., M.W, R.B., C.P.C, S.A.S., Y.C. and L.D. conducted experiments and acquired data. M.J., M.W, D.B. and J.N.Z. contributed to study design. M.J., M.W., R.B. and B.J.L analyzed and interpreted the data. M.W. Y.C., M.K., R.C.W., H.L.T., K.K., H.E.E., M.R.T., C.C.W., C.B., B.A.B., S.P. and J.M.M. performed recruitment, diagnostics, specimen collection and database maintenance., M.J., M.W., R.B., D.B. and J.N.Z. contributed to writing the manuscript.

## Conflict of Interest

The authors declare that there is no conflict of interest.

## Data Availability

The RNA-seq data discussed in of this publication have been deposited in NCBI’s Gene Expression Omnibus and are accessible through GEO Series accession number GSE212584.

The rest of datasets generated during and/or analyzed during the current study are available from the corresponding author on reasonable request.

## Notes

### Competing Interest Statement

The authors have declared no competing interest.

