## Supplemental Figures for "T-cell cellular stress and reticulocyte signatures, but not loss of naïve T lymphocytes, characterize severe COVID-19 in older adults": severity supplement.pdf

| Marker | Fluorochrome | Manufacturer |
|---|---|---|
| Live/dead | Blue | Thermofisher |
| CD3 | BV510 | Biolegend |
| CD4 | SparkBlue550 | Biolegend |
| CD8 | BUV395 | Thermo |
| CD95 | BV421 | Biolegend |
| CD28 | PEdazzle594 | Biolegend |
| CD45RA | BV570 | Biolegend |
| CCR7 | PerCP- | Biolegend |
| CD57 | BV605 | Biolegend |
| CD161 | BUV737 | BD |
| TCRγδ | PE-Cy5 | Thermo |
| CD19 | Spark NIR™ | Biolegend |
| CD20 | PerCP | Biolegend |
| HLA-DR | BUV805 | BD |
| CD127 | BV750 | BD |
| CXCR3 | BUV563 | BD |
| CD27 | BV785 | Biolegend |
| CD38 | BV650 | Biolegend |
| IgD | BV480 | BD |
| CD24 | BUV661 | BD |
| CD25 | BV711 | Biolegend |
| PD-1 | BB515 | BD |
| CXCR5 | BUV496 | BD |
| CD138 | SuperBright436 | Thermofisher |
| IgM | APC | Biolegend |
| Ki67 | PE | Biolegend |
| FoxP3 | AF700 | Biolegend |
| Perforin | APCFire750 | Biolegend |
| Eomes | e660 | Thermo |
| Tbet | PECy7 | Biolegend |
| Tigit | PerCPCy5.5 | Biolegend |

**Supplemental Table 1.** List of antibodies used for flow cytometric analysis

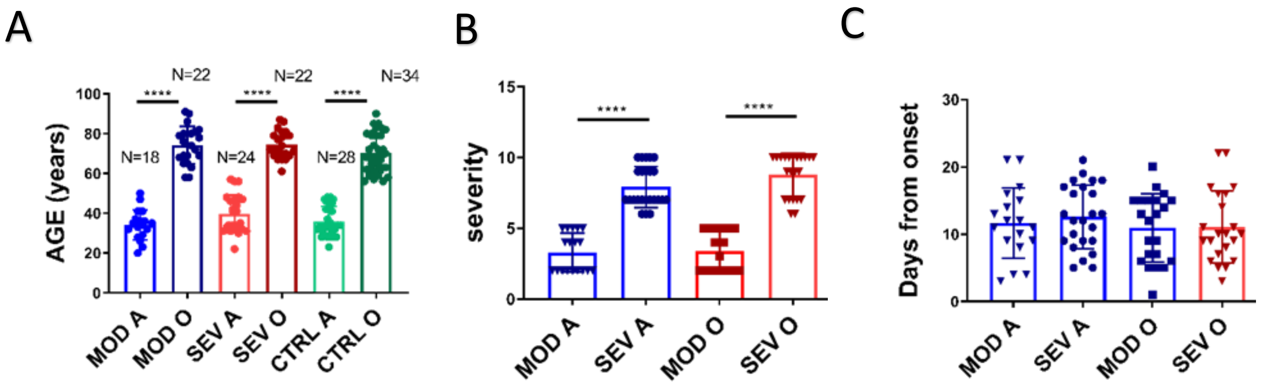


**Supplemental Figure 1. A**) 86 human subjects confirmed infected with SARS-COV-2 by PCR and compared to 62 naïve controls. N=18 adults with moderate disease (MOD A), N=22 older adults with moderate disease (MOD O), N=24 adults with severe disease (SEV A) and N=22 (SEV O) older adults with severe disease. N=28 adult controls (CTRL A). N=34 older adult controls (CTRL O). **B)** SARS-COV-2 positive participants were assigned to severe or moderate groups according to WHO Clinical Progression Scale. **C)** There was no difference between groups in time from onset of disease symptoms

A

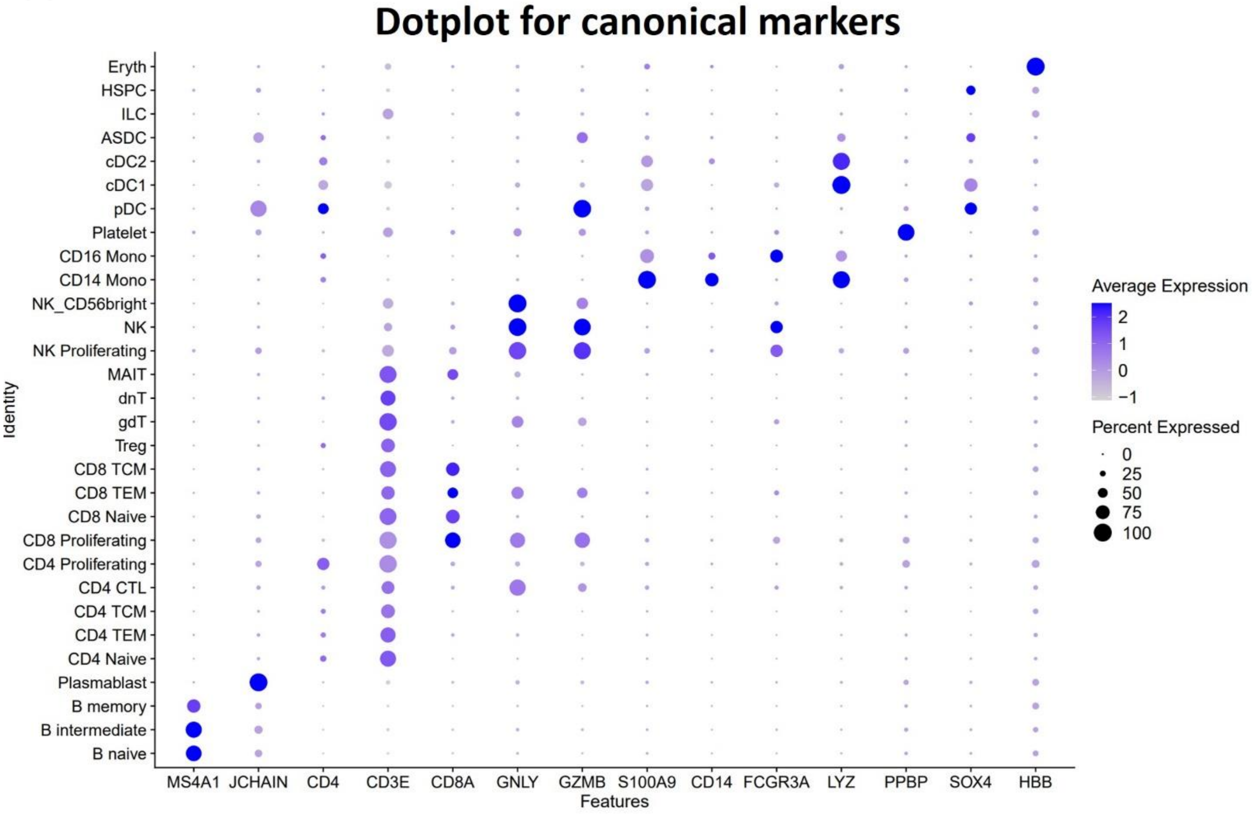


B

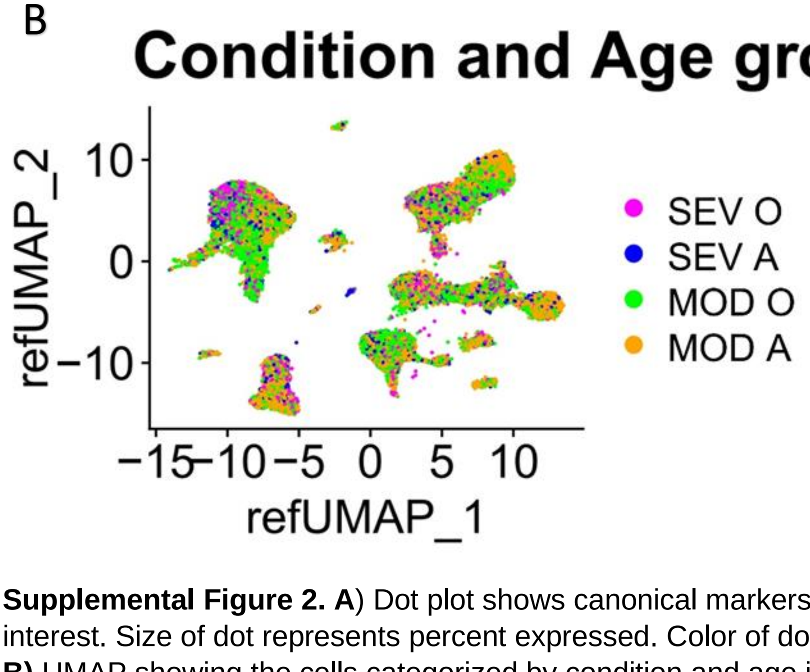


**Supplemental Figure 2. A**) Dot plot shows canonical markers for the different cell types of interest. Size of dot represents percent expressed. Color of dot represents average expression. **B)** UMAP showing the cells categorized by condition and age into four groups: SEV O, SEV A, MOD O and MOD A.

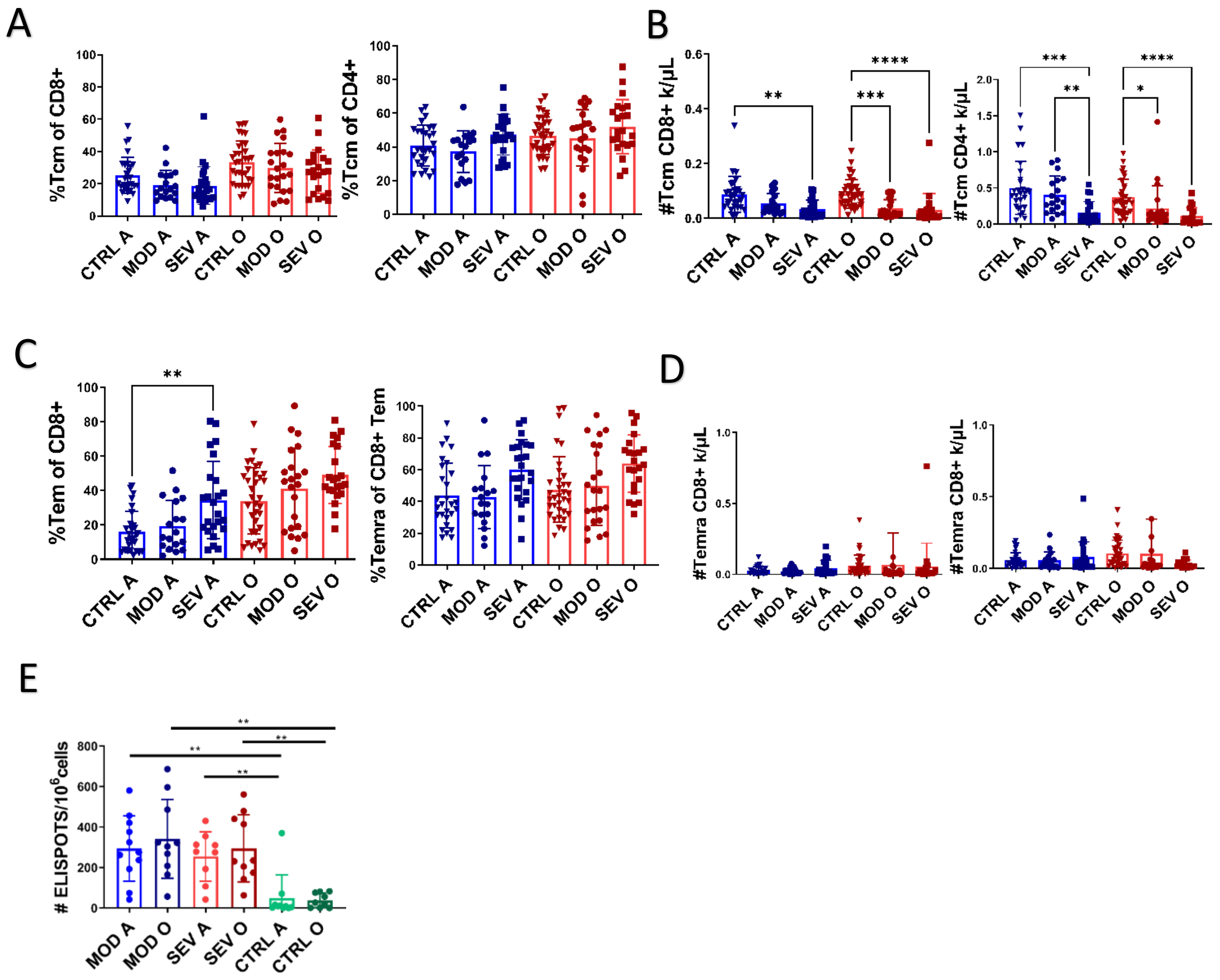


**Supplemental Figure 3. A)** No difference in percentage of central memory T cells between the groups. **B)** Decreased absolute counts central memory T cells in both severe groups **C)** Increased percentage of Tem CD8+ cells in adult severe COVID-19 participants. **D)** No difference between the groups in absolute counts of Tem cells. **E)** No difference in number of N, M, S antigen specific T cells between groups measured by ELISpot. Kruskal-Wallis test with Dunn's post hoc correction. Data presented as mean ± standard deviation. For all statistical differences*p<0.05, **p<0.01, ***p<0.001. ****p<0.0001.

A

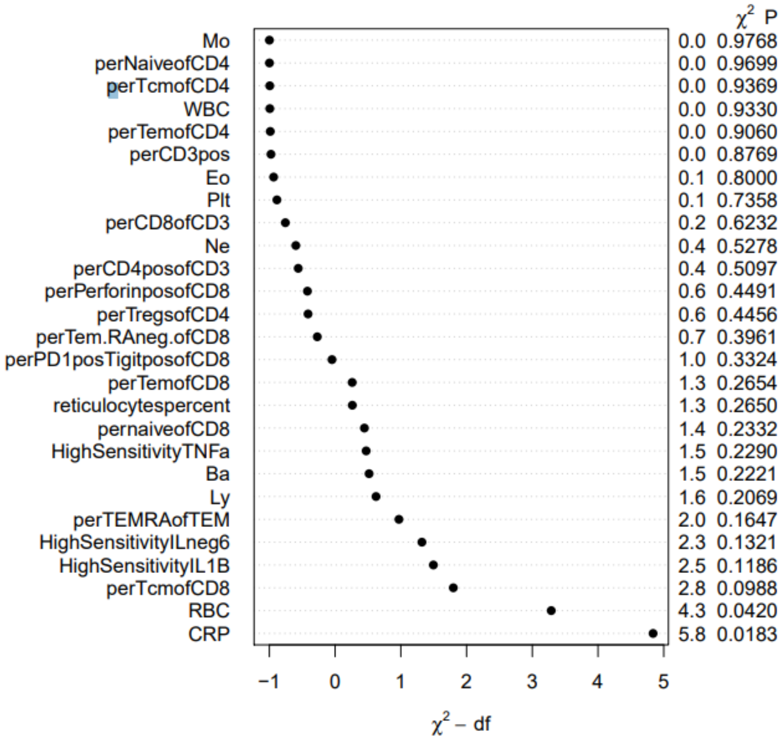


B

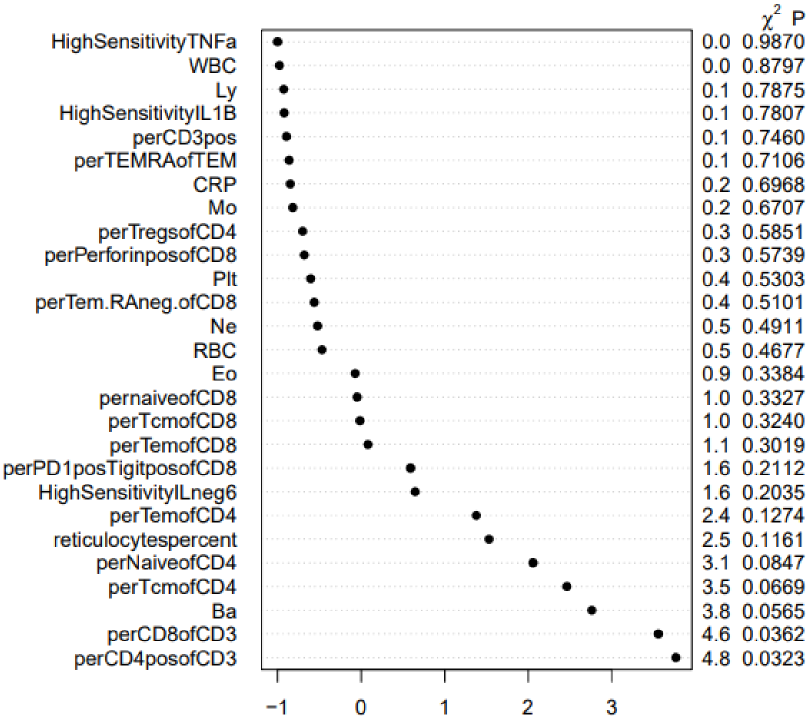


**Supplemental Figure 4.** Multivariable linear regression to predict dependence of **A)** severity and **B)** days from onset of symptoms to all predictor variables

Supplemental Figure 4

A

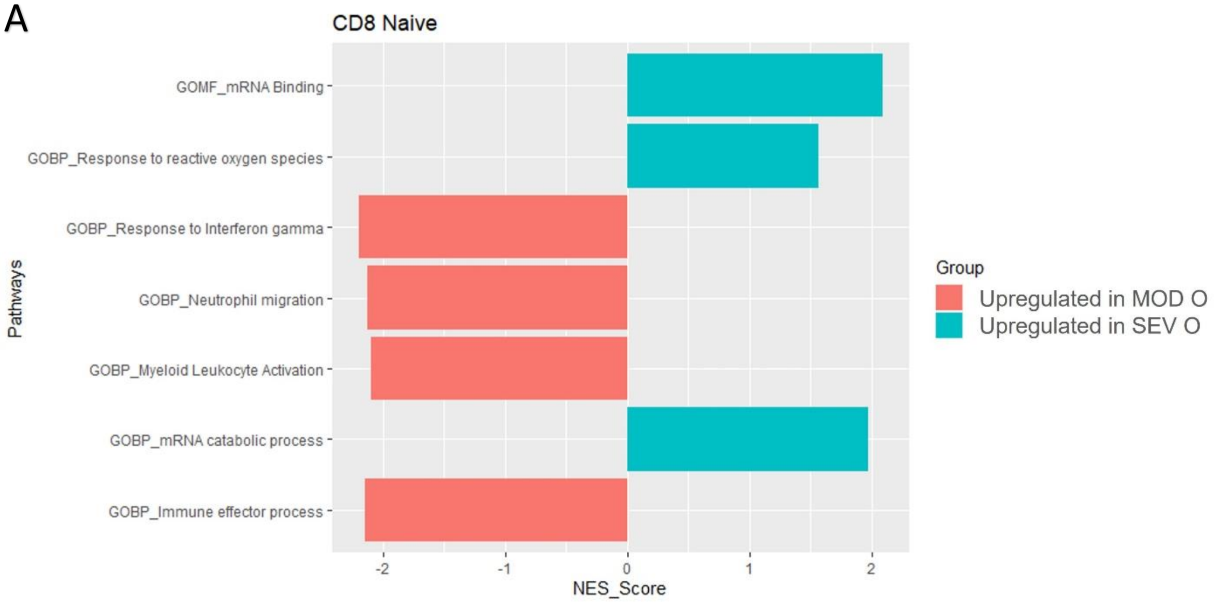
CD8 Naive
GOMF_mRNA Binding
GOBP_Response to reactive oxygen species
GOBP_Response to Interferon gamma
GOBP_Neutrophil migration
GOBP_Myeloid Leukocyte Activation
GOBP_mRNA catabolic process
GOBP_Immune effector process
Pathways
-2
-1
0
1
2
NES_Score
Group
Upregulated in MOD O
Upregulated in SEV O

B

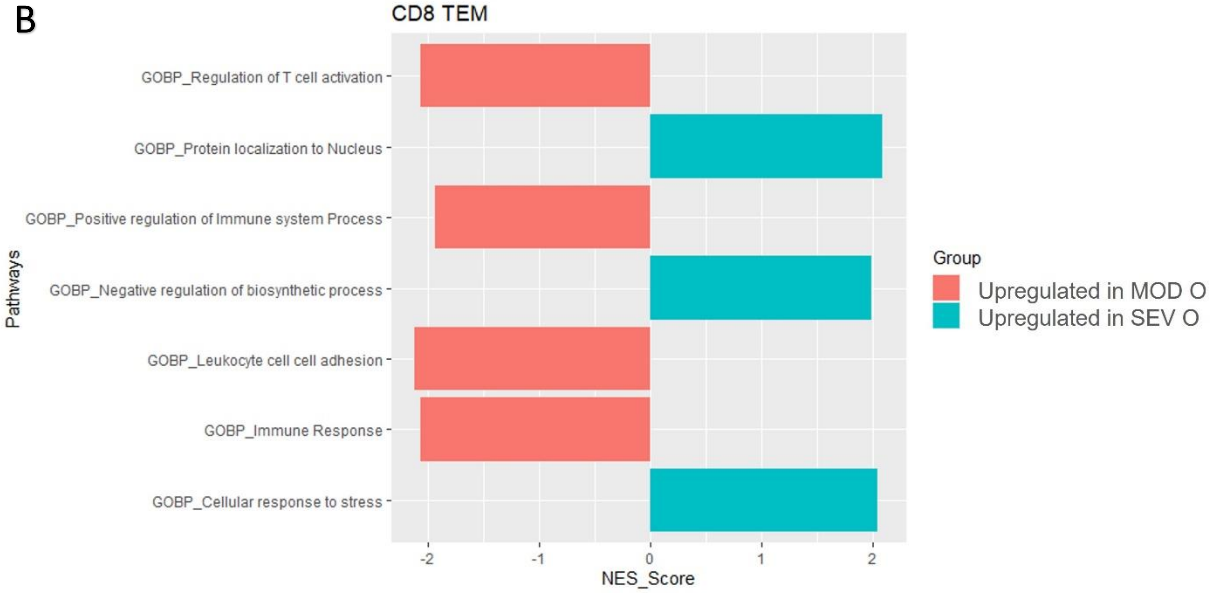
CD8 TEM
GOBP_Regulation of T cell activation
GOBP_Protein localization to Nucleus
GOBP_Positive regulation of Immune system Process
GOBP_Negative regulation of biosynthetic process
GOBP_Leukocyte cell cell adhesion
GOBP_Immune Response
GOBP_Cellular response to stress
Pathways
-2
-1
0
1
2
NES_Score
Group
Upregulated in MOD O
Upregulated in SEV O

C

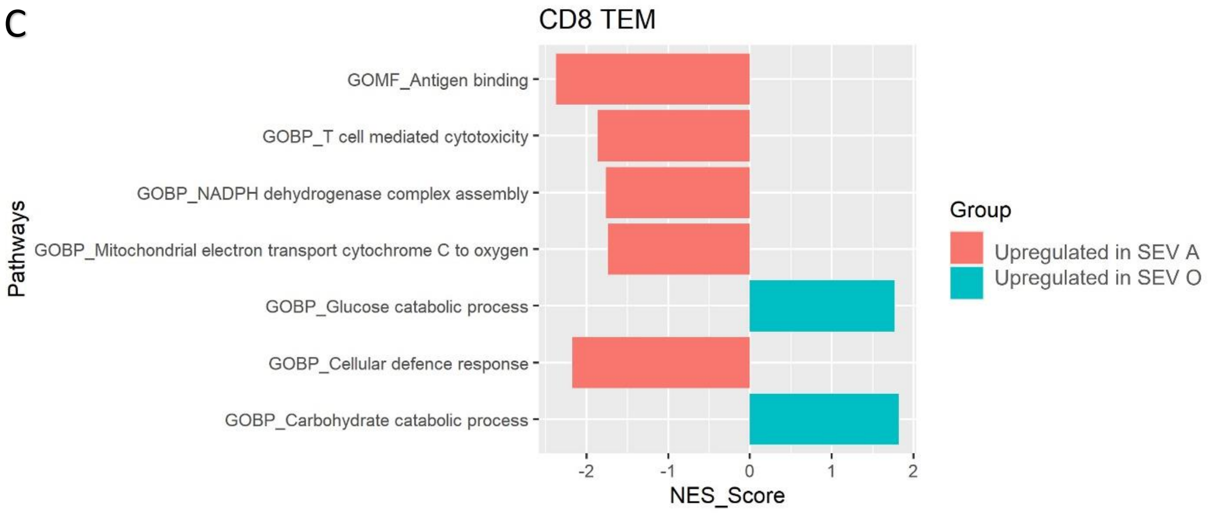
CD8 TEM
GOMF_Antigen binding
GOBP_T cell mediated cytotoxicity
GOBP_NADPH dehydrogenase complex assembly
GOBP_Mitochondrial electron transport cytochrome C to oxygen
GOBP_Glucose catabolic process
GOBP_Cellular defence response
GOBP_Carbohydrate catabolic process
Pathways
-2
-1
0
1
2
NES_Score
Group
Upregulated in SEV A
Upregulated in SEV O

D

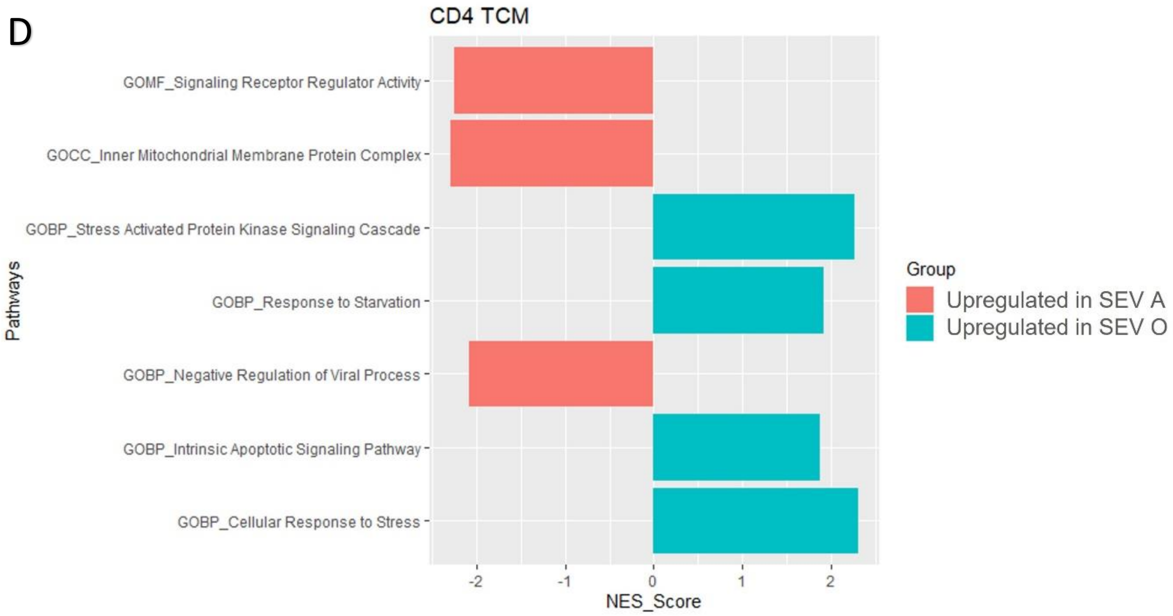
CD4 TCM
GOMF_Signaling Receptor Regulator Activity
GOCC_Inner Mitochondrial Membrane Protein Complex
GOBP_Stress Activated Protein Kinase Signaling Cascade
GOBP_Response to Starvation
GOBP_Negative Regulation of Viral Process
GOBP_Intrinsic Apoptotic Signaling Pathway
GOBP_Cellular Response to Stress
Pathways
-2
-1
0
1
2
NES_Score
Group
Upregulated in SEV A
Upregulated in SEV O

E

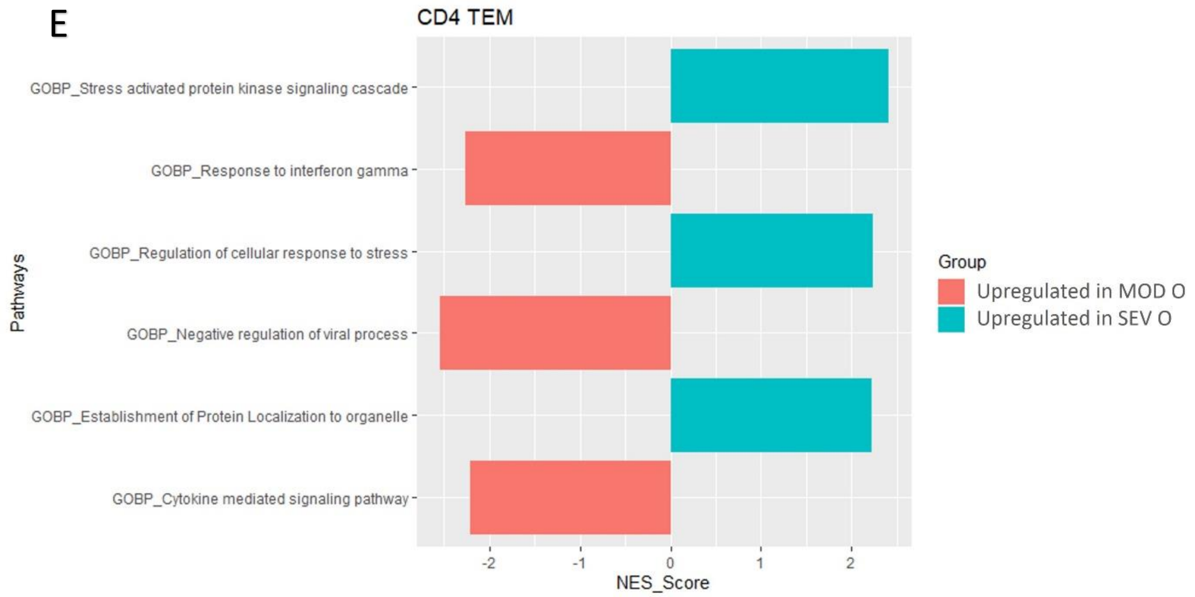
CD4 TEM
GOBP_Stress activated protein kinase signaling cascade
GOBP_Response to interferon gamma
GOBP_Regulation of cellular response to stress
GOBP_Negative regulation of viral process
GOBP_Establishment of Protein Localization to organelle
GOBP_Cytokine mediated signaling pathway
Pathways
-2
-1
0
1
2
NES_Score
Group
Upregulated in MOD O
Upregulated in SEV O

F

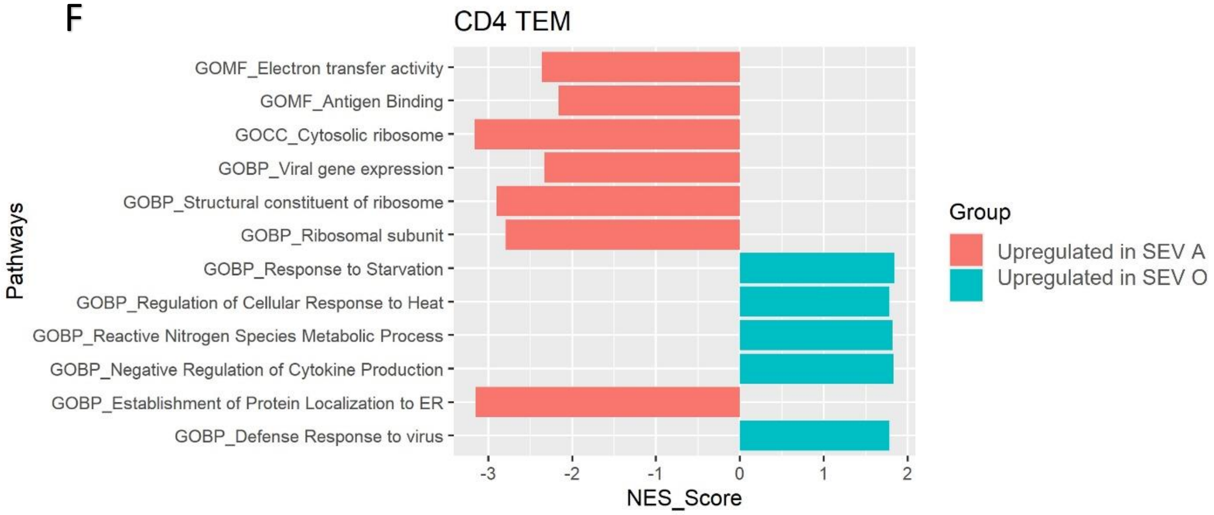
CD4 TEM
GOMF_Electron transfer activity
GOMF_Antigen Binding
GOCC_Cytosolic ribosome
GOBP_Viral gene expression
GOBP_Structural constituent of ribosome
GOBP_Ribosomal subunit
GOBP_Response to Starvation
GOBP_Regulation of Cellular Response to Heat
GOBP_Reactive Nitrogen Species Metabolic Process
GOBP_Negative Regulation of Cytokine Production
GOBP_Establishment of Protein Localization to ER
GOBP_Defense Response to virus
Pathways
-3
-2
-1
0
1
2
NES_Score
Group
Upregulated in SEV A
Upregulated in SEV O

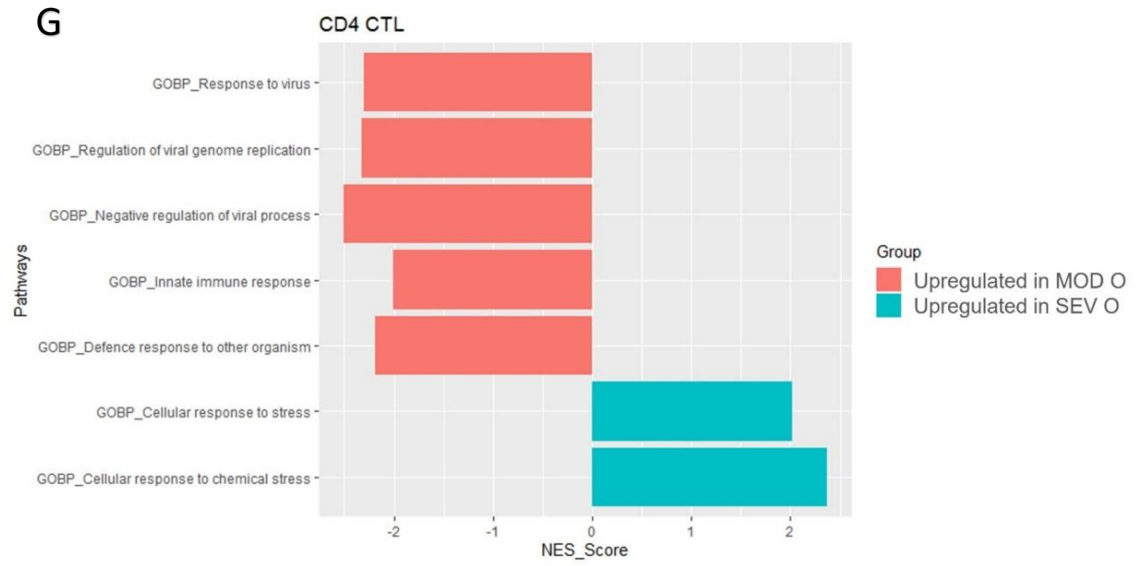


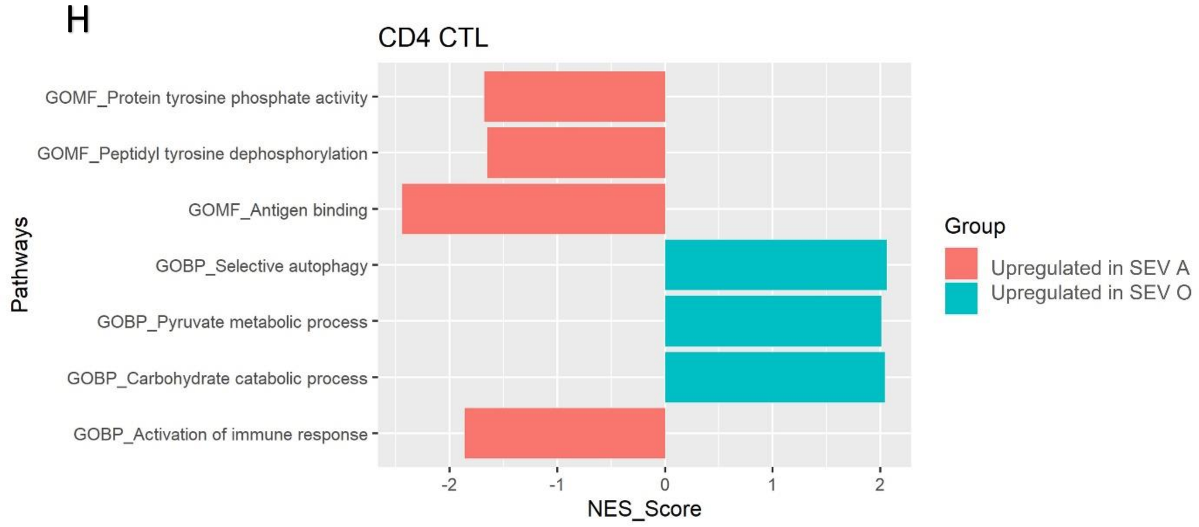


**Supplemental Figure 5.** Barplots showing differential enrichment of significant (FDR≤0.18) pathways in the various subsets of the CD4 and CD8 populations based on their normalized enrichmnt score (NES) calculated based on their differential gene expressions (adjusted p value<0.00001). **A)** CD8 Naïve cells between MOD O and SEV O, **B)** CD8 TEM cells between MOD O and SEV O, **C)** CD8 TEM cells between SEV A and SEV O, **D)** CD4 TCM cells between SEV A and SEV O, **E)** CD4 TEM cells between MOD O and SEV O, **F)** CD4 TEM cells between SEV Aand SEV O, **G)** CD4 CTL cells between MOD O and SEV O and **H)** CD4 CTL cells between SEV A and SEV O. Pathway enrichment analysis was performed using the GSEA tool against the Hallmark gene set “c5.go.v7.4.symbols.gmt[Gene Ontology]” of the MSigDB.

**A**

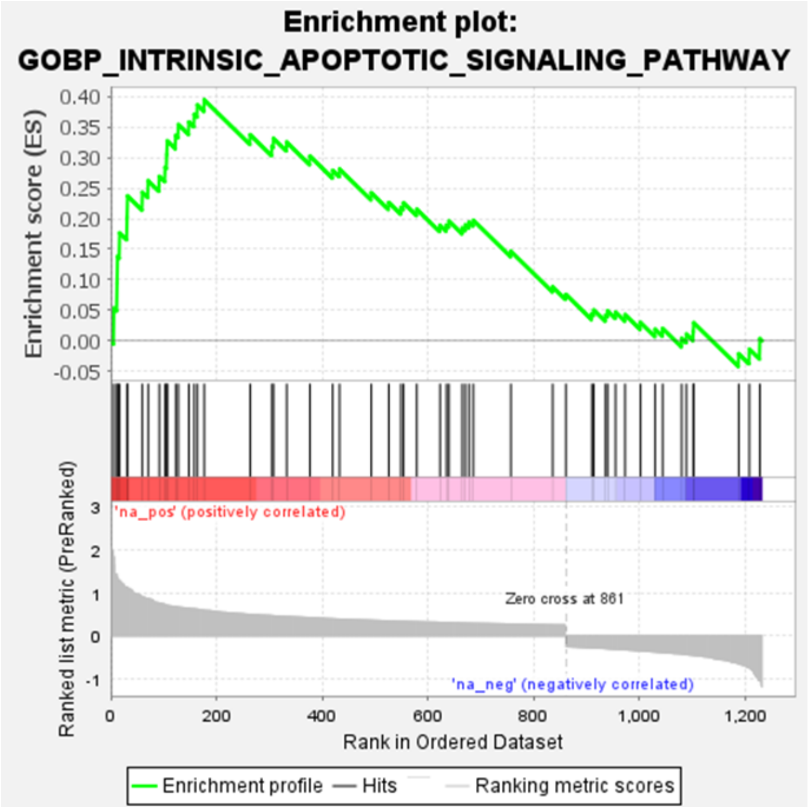


**B**

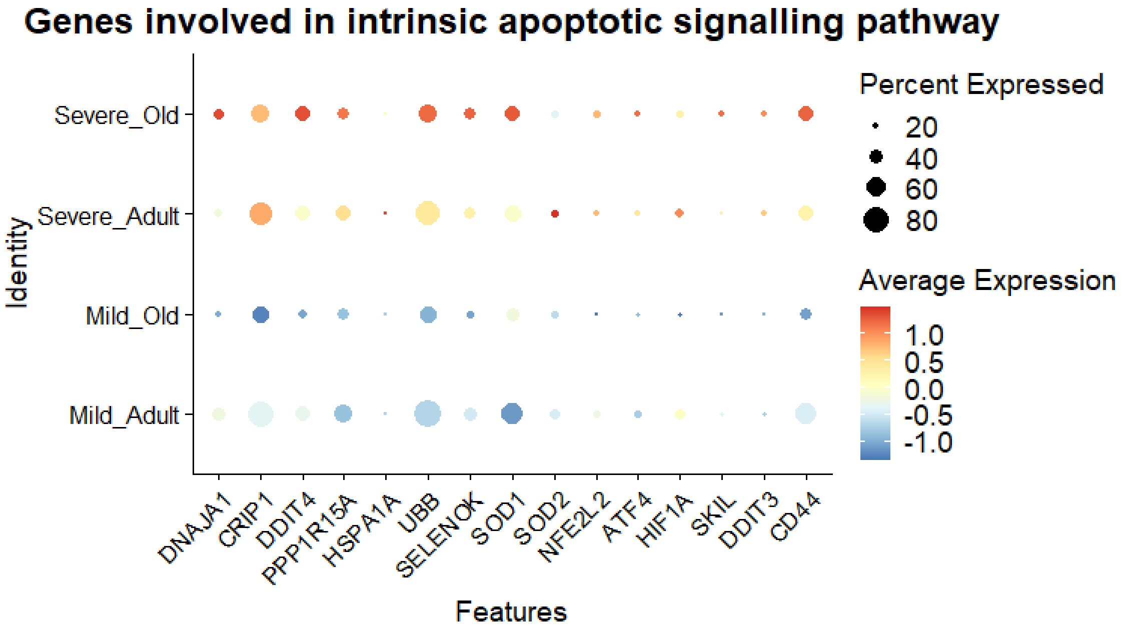


**Supplemental Figure 6. A)** GSEA enrichment profile of 'Intrinsic Apoptotic Signaling pathway' with enrichment score and **B)** Expression levels of genes involved in 'Intrinsic Apoptotic Signaling pathway' as per SEV O, SEV A, MOD A and MOD O categories. Size of dot represents percent expression and color gradient represents average expression.
